# WAT’s up!? – Organ-on-a-chip integrating human mature white adipose tissues for mechanistic research and pharmaceutical applications

**DOI:** 10.1101/585141

**Authors:** Julia Rogal, Carina Binder, Elena Kromidas, Christopher Probst, Stefan Schneider, Katja Schenke-Layland, Peter Loskill

## Abstract

Obesity and its numerous adverse health consequences have taken on global, pandemic proportions. White adipose tissue (WAT) – a key contributor in many metabolic diseases – contributes about one fourth of a healthy human’s body mass. Despite its significance, many WAT-related pathophysiogical mechanisms in humans are still not understood, largely due to the reliance on non-human animal models. In recent years, Organ-on-a-chip (OoC) platforms have developed into promising alternatives for animal models; these systems integrate engineered human tissues into physiological microenvironment supplied by a vasculature-like microfluidic perfusion. Here, we report the development of a novel OoC that integrates functional mature human WAT. The WAT-on-a-chip is a multilayer device that features tissue chambers tailored specifically for the maintenance of 3D tissues based on human primary adipocytes, with supporting nourishment provided through perfused media channels. The platform’s capability to maintain long-term viability and functionality of WAT was confirmed by real-time monitoring of fatty acid uptake, by quantification of metabolite release into the effluent media as well as by an intact responsiveness to a therapeutic compound. The novel system provides a promising tool for wide-ranging applications in mechanistic research of WAT-related biology, in studying of pathophysiological mechanisms in obesity and diabetes, and in R&D of pharmaceutical industry.

## Introduction

The global obesity pandemic poses one of today’s biggest challenges to public health. Each year, 2.8 million people die from causes related to overweight or obesity.(1) Since the 1970s, the worldwide prevalence of obesity has nearly tripled, also leading to an upsurge in associated comorbidities such as type 2 diabetes (**figure 1a**).(2) Future projections reveal the gravity of this public health crisis: the prevalence rates of childhood obesity are increasing at an alarming pace(3), and by 2030, more than 50% of U.S. adults are predicted to be obese.(4) White adipose tissue (WAT) is the principal organ in obesity. In healthy human adults, WAT comprises approximately 20-25% of the total body mass, thus constituting the second largest organ, after the skin. In obese individuals, WAT’s contribution to the total body mass may become as high as 50% (**figure 1b**).(5)

**Fig. 1.**
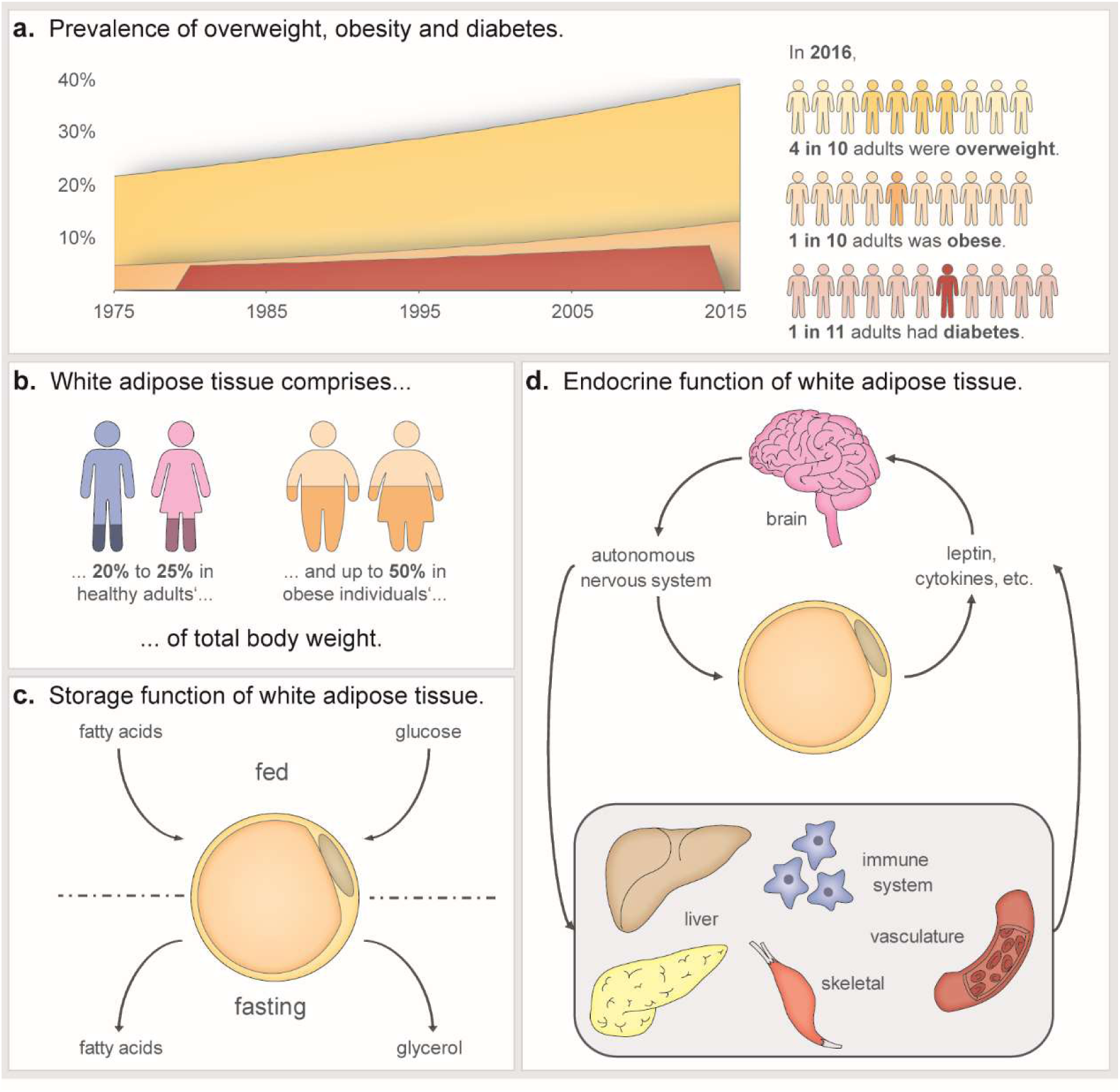
Relevance of research on WAT. **a.** The worldwide prevalence of obesity has nearly tripled since 1975; in 2016, almost 40% of adults aged 18 and over could be classified as overweight, 13% of them were even obese. This developments has coincided with a rising prevalence of diabetes, which in 2016 affected approximately 9% of adults worldwide. T2DM – the most prevalent form of diabetes – often develops during chronic positive energy balance, e.g., by a combination of excessive energy intake and physical inactivity.(1,2) **b.** WAT’s contribution to body mass is 20-25% in healthy individuals, and up to 50% in the obese.(5) **c.** Traditional view of WAT as an organ primarily for energy storage. **d.** Additional, modern-day concepts of WAT functionality, including extensive endocrine functions.

WAT is tightly involved in the two most important functions of an organism – energy homeostasis and reproduction.(6) In energy homeostasis, not only does WAT act as the main storage site of excess dietary energy (**figure 1c**), it also performs crucial endocrine and metabolic functions (**figure 1d**).(7,8) WAT can sense the body’s energy status, and respond appropriately, either by storing fuel, in the form of triacylglycerides, or by releasing it as glycerol and fatty acids, for ultimate delivery to organs in need. While this classically described role of WAT already entailed extensive crosstalk between WAT and other organs, its inter-organ communications extend beyond simple feedback loops activated by fed-or fasted states. Endocrine functions of white adipocytes, and other WAT-resident cells in the stromal vascular fraction, are performed by the release of a variety of adipokines (adipose-associated cytokines) which affect the functioning of the brain, liver, pancreas and immune system.(9) Besides managing nutritional homeostasis, WAT contributes to the regulation of the hypothalamic-pituitary-gonadal axis by secreting and metabolizing sex steroids.(7) Especially adipose tissue intracrinology, i.e., the modulation of sex steroid levels by significant numbers of locally expressed enzymes that activate, convert, or inactivate circulating steroid hormones, plays important roles in human reproductive function.(7,10,11) Given its prominent endocrine functions and extensive cross-talk with other organs, it comes as little surprise that abnormal amounts, or altered functioning of WAT may result in wide-ranging disorders, including hepatic and cardiovascular diseases,(12,13) diabetes and cancer.(14–16)

In line with its important roles in metabolism, inflammation and cancer, WAT has emerged as a drug target with major therapeutic potential for a variety of diseases.(17–19) Additionally, storage metabolism by WAT can have a major impact on the efficacy of drug therapies, such as those for cardiovascular diseases and cancer.(20,21) For instance, adipocytes may metabolize and inactivate the chemotherapeutic, Daunorubicin,(21) which strongly affects the efficacy of this anticancer therapeutic. Furthermore, the capacity of WAT to sequester hydrophobic compounds, gives the tissue a prominent role in absorption, distribution, metabolism, and excretion (ADME) processes.

Even though WAT plays such a significant role in many diseases, surprisingly little is known about pathophysiological processes of WAT, especially when considering the ever-increasing prevalence of obesity and its co-morbidities. An important reason for this lacking insight is that mechanistic studies in humans often involve unacceptable health risks. Hence, most research depends on clinical observation, genome-wide association studies (GWAS) as well as animal studies, all with limited success.(22) Although animal models have led to many insights in obesity and diabetes, they often are lacking in predictive validity for human body functioning, first because of important species differences in nutrition and metabolism, and second because the distinct, unique physiological and pathophysiological roles of WAT in humans(23). Alternatively, conventional, cell culture-based *in vitro* models have been widely utilized and are significantly less controversial ethically. Typically, researchers have used a variety of cell sources, ranging from (immortalized) murine, to primary human (pre-)adipocytes, each featuring distinct advantages and limitations.(24) *In vitro* differentiation of adipogenic progenitor cells, i.e., either pre-adipocytes or multipotent stem cell lines, or (induced) pluripotent stem cells, is often the method of choice due to the availability of donor-specific information, and the capacity of these cells to undergo expansion and cryopreservation while retaining their characteristic fat depot. Yet, the *in vitro* use of differentiated adipocytes has two major drawbacks: unlike their *in situ* matured congeners, (i) their lipid contents never reach a state of unilocularity, but remain distributed among multiple small lipid vacuoles, and (ii) their secretome differs immensely in the relative proportions of adipose-associated hormones(25,26). Both phenomena reflect a pre-mature state of the in vitro differentiated adipocytes, which strongly suggests that mature human adipocytes provide the best recapitulation of a mature human adipocyte physiology. Yet, the **ex vivo** culture of primary human adipocytes is extremely challenging, due to issues with the cells’ buoyancy, fragility and de-differentiation, which so far have hindered development of robust protocols for long-term culturing. Additionally, inter-individual variability complicates the interpretation of study results based on samples from different donors.

Overall, it is of upmost importance to develop microscale platforms that provide microphysiological environments for the long-term culture of white adipocytes in structures that may recapitulate *in vivo* physiology, and functionality based on a minimal amount of cells. By combining modern techniques in microfabrication, biomaterials and tissue engineering, organ-on-a-chip technology has enabled the construction of promising platforms for mechanistic and pharmaceutical studies, that have great potential for disease modeling as well as the optimization of personalized medical treatments of obesity and diabetes.(27–30) The integration of tissues with *in vivo*-like structure and functionality in perfused microenvironments is of particular interest when studying endocrine tissues and multi-factorial diseases, due to the possibility to combine individual chips into multi-organ systems.(31,32) However, although organ-on-a-chip research has burgeoned in recent years, and numerous platforms have been developed for many organs and tissues, WAT appears to have been largely overlooked, and only a few relevant efforts have been undertaken.(25) Several research groups injected pre-adipocytes from murine(33,34) or human(35,36) sources into microfluidic chambers, and were able to subsequently induce adipogenesis. Others directly introduced primary adipocytes from mice into mesoscopic, perfused reservoirs.(37) All of these innovative approaches, however, did not achieve the goal of generating human white adipose tissue structures with *in vivo*-like physiology.

Here, we present the first OoC platform that integrates human primary mature adipocytes into a perfused microfluidic chip. The human WAT-on-a-chip consists of multiple, tissue-specific chambers that are fluidically connected to a vasculature-like microchannel, while being shielded from the shear forces of the perfusion fluid. Using specifically tailored isolation and injection protocols, 3D microtissues based on freshly isolated adipocytes are generated inside a series of individual chambers. Thus, a large number of independent replicate cell cultures from individual donors can be produced and kept viable – as well as functional – for over one month. By analyzing media effluents and taking advantage of the optical accessibility of the tissue chambers, the patency of cells as well as key cell-physiological aspects, e.g., fatty acid metabolism and drug responsiveness, could be monitored successfully.

## Results and Discussion

### Concept and characterization of the microfluidic platform

In order to generate a microphysiological environment capable of integrating human WAT, and maintaining tissue viability and functionality during long-term culture, we developed a specifically tailored microfluidic platform featuring a footprint comparable to the size of a stamp (2 x 2 cm). The essential features of the platform, depicted in **figure 2a**, are 3D-tissue chambers and perfusable microchannels that are separated by an isoporous membrane. Each chip platform houses two identical, independent systems each featuring eight individual tissue chambers. All tissue chambers are located at the end of individual branches of a main channel, which can be accessed via a common inlet for cell injection. The tissue chambers are specifically designed to accommodate adipose tissue, particularly its large-sized, fragile and buoyant adipocytes: they are cylindrical structures with a diameter of 720 µm and a height of 200 µm. The tissue-chamber microstructures are encased by transparent glass (cover slip or microscope slide) at the bottom, enabling easy visual inspection of the tissues with most current types of microscope, and by an isoporous membrane on the top side. The membrane was specifically functionalized (**supplement S2**) to ensure a tight sealing of chip components, and to provide a porous barrier to the adjacent media microchannels. The channels can be connected to external devices (e.g. syringe-or peristaltic pumps), to achieve a vasculature-like perfusion with media. This allows for a precisely controllable convective transport of nutrients, metabolites, and other dissolved molecules towards, as well as away from, the tissue chambers, mimicking the *in vivo* circulation of blood. It also opens up the possibility to administer compounds with high temporal resolution. The membrane further ensures that convective transport is restricted to the media channels, thereby shielding the tissue chambers from non-physiological shear forces (**figure 2b**). Through the micropores, dissolved molecules may diffuse quickly in and out of the tissue chambers, as confirmed by computational modeling and by dynamic tracking of a fluorescent dye in different channel-and chamber locations (**figure 2b**). In sum, the membrane constitutes an endothelial-like barrier that separates perfused media from the chip-embedded tissues. Although this artificial barrier admittedly does not recapitulate active transport processes occurring *in vivo*, it does provide a potential scaffold for the inclusion of endothelial cells in future generations of the platform.

**Fig. 2.**
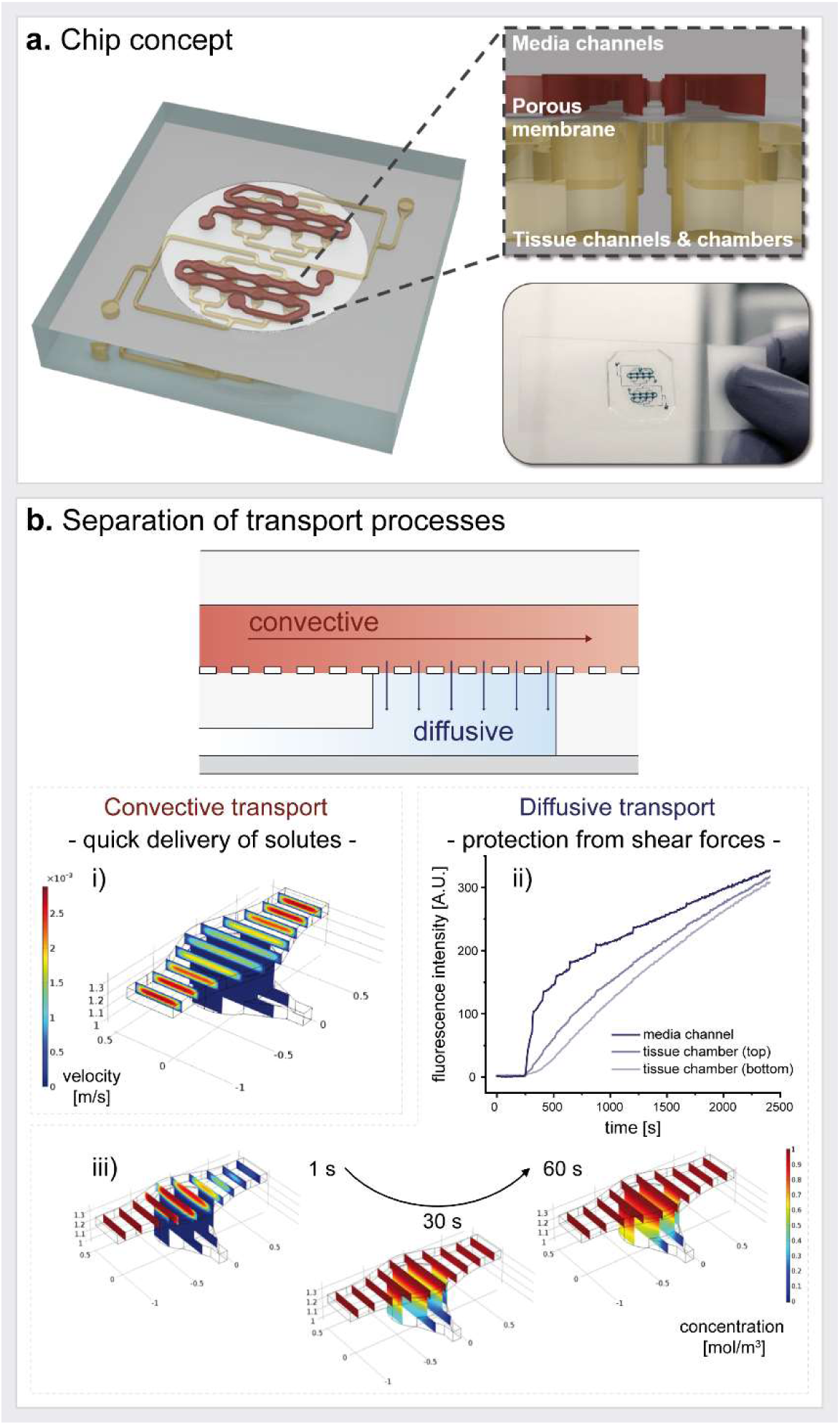
Concept of the WAT-microphysiological platform and microfluidic perfusion. **a.** Schematic of the chip platform featuring two independent systems with eight tissue chambers each. As shown in the cross-sectional zoom-in, tissue chambers, which feature tissue channel inlets at the bottom, are located right below the perfusable media channels, and separated from them by a microporous membrane. **b.** During flow, the chip’s circuitry imposes a separation of the two transport processes, convection and diffusion. Convective flow is confined to the vasculature-like media channels, as confirmed by computational modeling (i). The tissue chambers are supplied by a diffusive transport as confirmed by observable diffusion of a fluorescent dye from the media channel into a hydrogel-filled tissue chamber (ii), as well as by computational modeling (iii).

### Generation and functional validation of human white adipose tissue on-chip

Two major difficulties of mature adipocytes – their buoyancy and fragility – have traditionally hampered the *in vitro* culture of WAT. We developed specifically optimized cell-isolation and - injection protocols to enable the integration of freshly isolated, mature primary adipocytes from humans into the tissue chambers of our microphysiological platform. By suspending the isolated cells into a collagen hydrogel before injection, their viability and integrity could be preserved. The use of the hydrogel ensures both the protection and the 3D arrangement of the fragile cells during the formation of the microtissue. Moreover, the hydrogel and the location of the cell-injection inlets on the bottom of the tissue chambers (**figure 2a, figure 3**) help to overcome the issue of adipocyte buoyancy. Successful cell loading generates 8 independent, parallel 3D WAT microtissues per system (and 16 per platform).

**Fig. 3.**
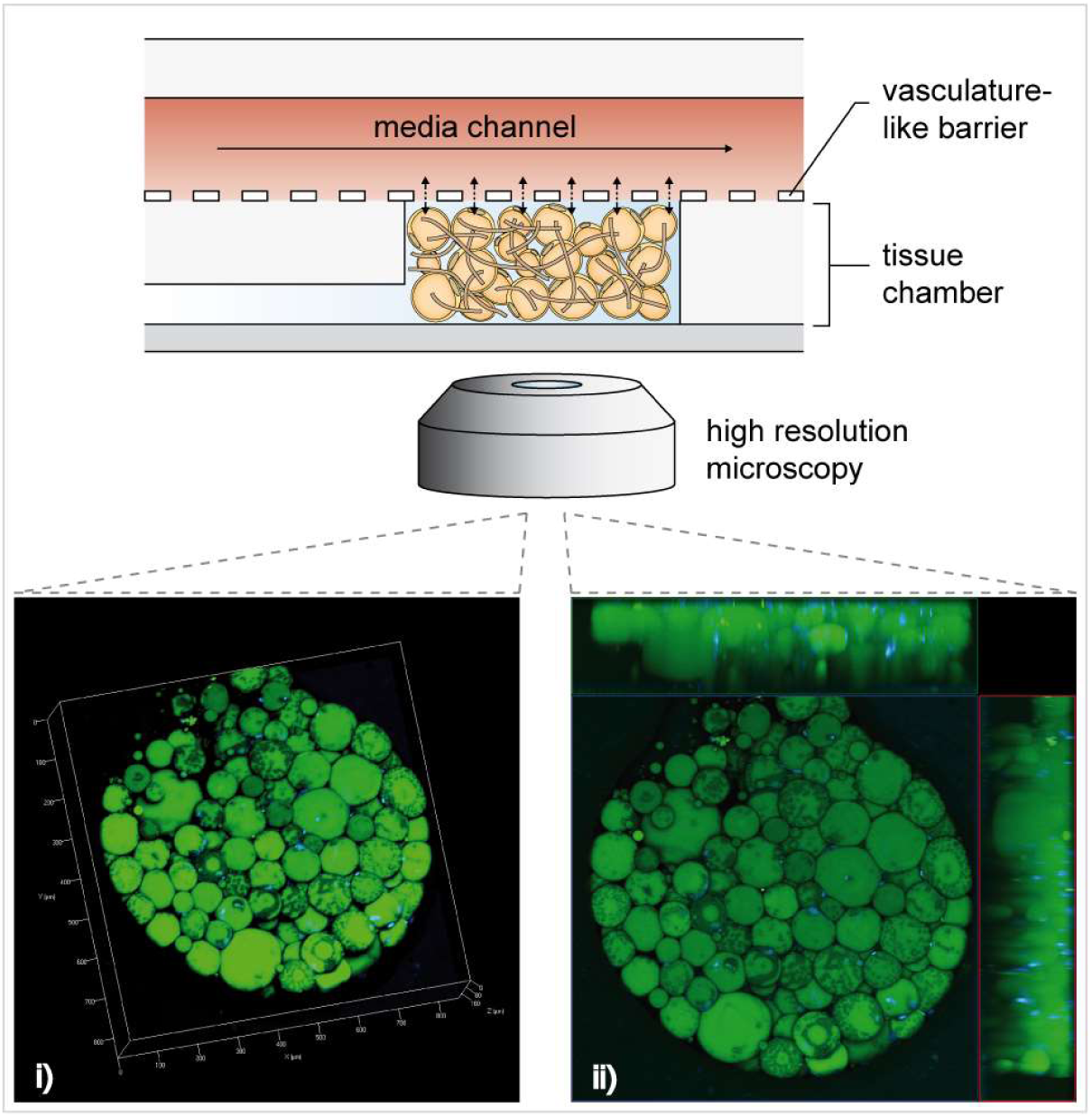
Structural characterization of human WAT-on-a-chip. On day 6 of on-chip culture, intracellular lipid vacuoles (green; neutral lipid stain BODIPY™ 493/503) and nuclei (blue; DAPI) were visualized by confocal imaging. As shown by a 3D-rendered Z-stack of one WAT-chamber (i) (steps on the scale in 100 µm) and the corresponding maximum intensity projection with two sectional planes (ii), the individual adipocytes were unilocular and formed a densely packed 3D microtissue throughout the tissue chamber.

Due to the microscale footprints of chambers and platform, only a small number of cells are required per chamber. This is of particular importance, as the tissues are based on primary, non-proliferating cells, and multiple chips (independent replicates) can be loaded with cells from a single biopsy, enabling testing under multiple experimental and control conditions of microtissues from the same donor. This is of crucial advantage when dealing with inter-donor variability, while it also enables patient-specific experiments.

### Structural analysis of WAT on-chip

To investigate the structural arrangement of the generated adipose tissue, we established a staining protocol that enabled the on-chip visualization of intracellular lipid vacuoles using a neutral lipid stain. After six days of on-chip culturing, confocal imaging confirmed the formation of dense, 3D tissues composed of large unilocular adipocytes (**figure 3**). The observable, mature phenotype of adipocytes suggests that the tissue-generation and -culture methods were capable of creating and maintaining physiological microenvironments and conditions. Inappropriate culture conditions have been frequently reported to induce cell dedifferentiation(38,39) which would lead to highly proliferative dedifferentiated fat cells (DFAT) with multilineage character.(40) One of the first hallmarks of dedifferentiation is the re-organization of lipids into multiple lipid droplets (i.e., a multilocular phenotype) and an increased rate of lipolysis.(39)

### Validation of WAT functionality on-chip

To assess the capability of the WAT-on-a-chip system to maintain viability and functionality of the integrated tissues, we deployed a comprehensive toolbox of chip-specific, functional readout methods. First, the viability and integrity of the vast majority of cells after eight days of on-chip culture was confirmed using a FDA/PI-based live/dead-staining protocol tailored specifically for the chip’s configuration (**figure 4a**). In addition, to verify that the objective of any microphysiological OoC system – the recapitulation of *in vivo* functionality – was indeed attained, we examined multiple functional endpoints. Taking advantage of the optical accessibility and continuous medium perfusion through our system, we performed non-invasive imaging and analysis of media effluents under several conditions.

**Fig. 4.**
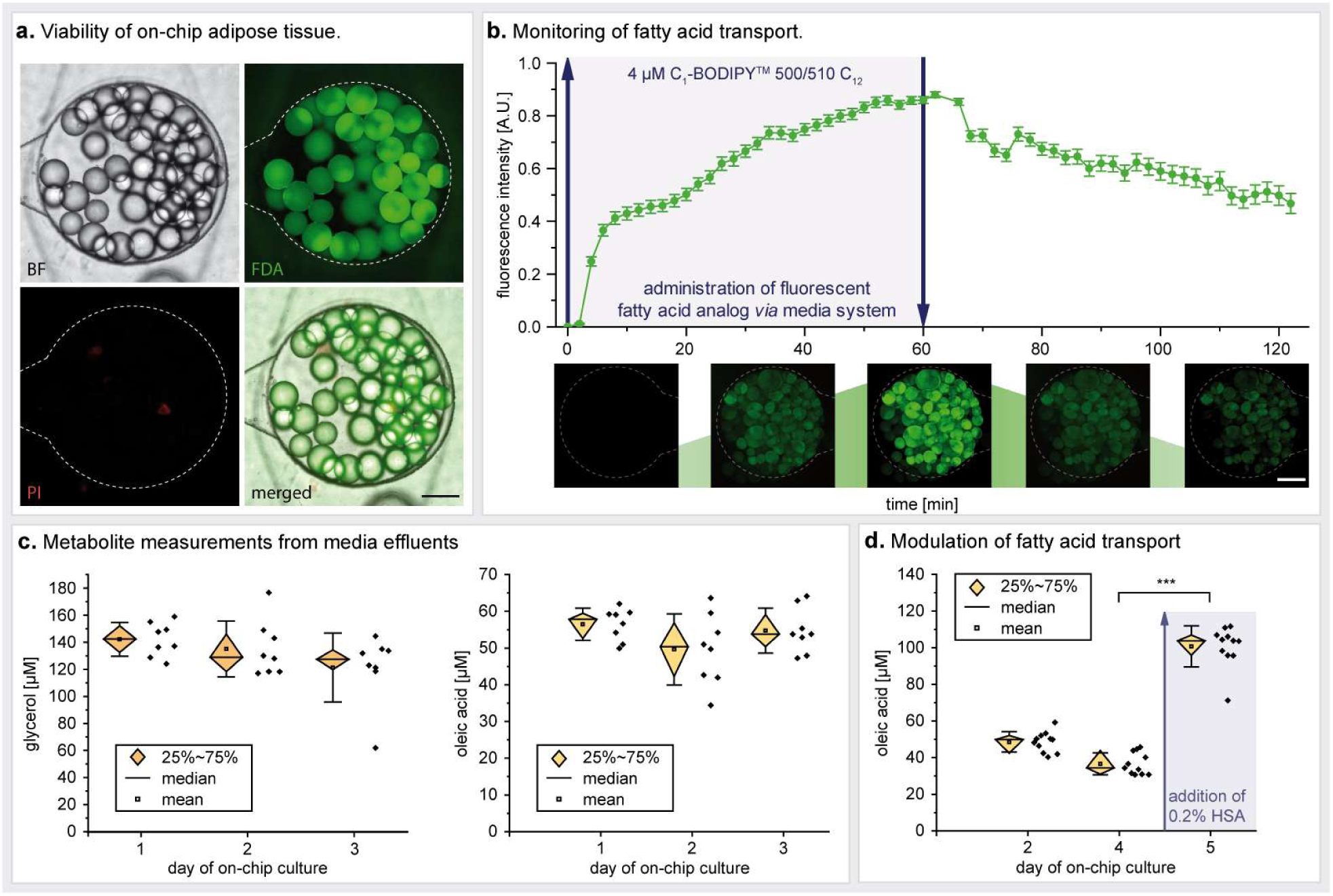
Functional validation of the human WAT-on-a-chip. **a.** Live/dead-staining of WAT on day 8 of on-chip culture. FDA (green) marks live adipocytes by confirming membrane integrity and esterase activity in the vast majority of the integrated adipocytes. PI (red) staining indicates a very low number of apoptotic/necrotic and dead adipocytes. Scale bar equals 150 µm. **b.** On-line monitoring of fatty acid transport on day 6. Monitoring of fatty acid uptake and accumulation was realized by the addition of a fluorescent fatty acid analog (4 µM BODIPY™ 500/510 C1, C12) to the perfused medium for one hour, followed by measurement of the mean fluorescence intensity. Upon removal of the fatty acid analog from the medium, fatty acid release could be observed (n = 47 individual tissue chambers). Scale bar equals 150 µm. **c.** Evaluation of fatty acid metabolites in media effluents. Using colorimetric assays, glycerol and oleic acid (representative of non-esterified fatty acids) concentrations were found to be stable for the first three days of on-chip culture (n = 8 systems with 8 tissue chambers each). **d.** Upon addition of 0.2% w/v human serum albumin (HSA) to the culture medium for 24 h, fatty acid release significantly increased compared to that during previous days (p < 0.005; n =11 systems with eight tissue chambers each).

To monitor fatty acid metabolism in real-time, we added the fluorescently tagged fatty dodecanoic acid (C_12_H_24_O_2_) analog (BODIPY™ 500/510 C1, C12) to the perfusion medium, and characterized the dynamics of fatty acid influx into, and efflux from, the WAT. This assay represents a powerful tool that allows non-invasive assessment of the cellular uptake of fatty acids, their accumulation in intracellular lipid vacuoles, and their release from adipocytes when the fatty-acid analog is removed from the medium. Utilizing time-resolved fluorescence microscopy, we were able to observe these characteristic uptake and secretion kinetics in the WAT cultured on-chip (**figure 4b**).

The continuous, convective transport by the perfusion flow, of metabolized and secreted factors away from the tissues, provides opportunities for effluent sampling in a time-resolved manner, allowing the dynamics of e.g. metabolism, secretion, and endocrine functionality to be measured. Using colorimetric assays, we characterized the levels of oleic acid, a representative of non-esterified fatty acids (NEFAs), and of glycerol in the effluent medium, and found that their concentrations remained very stable for several days of culturing (**figure 4c**). The measured NEFA and glycerol levels were comparable to levels from human subcutaneous adipose tissue explants immediately after biopsy(41). In a similar approach, we examined the impact on fatty acid release of a 0.2 (w/v) % supplementation of the perfusion medium with human serum albumin (HSA). After 24 h of perfusion, we observed a significant increase of NEFA levels in the effluent (**figure 4d**). Thus, in our system, effluent sampling provided a powerful, non-invasively obtained readout of secretome and metabolome dynamics with high temporal resolution (depending on the sensitivity of the employed assay). This constitutes an effective link for the cross-correlation with clinical data and biomarker development.

### Long-term functionality of integrated white adipose tissue

One of the main challenges of conventional adipocyte *in vitro* culture is the long-term maintenance of functional stability. As discussed above, issues related to buoyancy, fragility and de-differentiation of adipocytes, typically limit the duration of integrity of *in vitro* cultures to a couple of days at most(41,42). Our structural and functional characterization of the WAT-on-a-chip indicated that this microphysiological platform is capable of maintaining stable culture conditions over prolonged time periods. However, to investigate the potential of the platform to support a robust, long-term culture, we performed fatty acid uptake assays on the tissues after 6 days, as well as after 36 days of on-chip culture (**figure 5**). We observed that even after more than a month of *in vitro* culture, adipocytes still showed functional uptake and accumulation of fatty acids. Interestingly, a difference in initial uptake rates emerged, which might be related to size limitation of the “long-term-fed” cells on day 36. Still, these findings highlight that the WAT-on-a-chip is indeed capable of maintaining the functionality of adipose tissue for much longer periods than are conventional methods. This brings within reach a variety of novel opportunities for *in vitro* studies, e.g. of long-term effects of nutrition, repeated exposure of potential toxicants, or of long-term endocrine dynamics.(43)

**Fig. 5.**
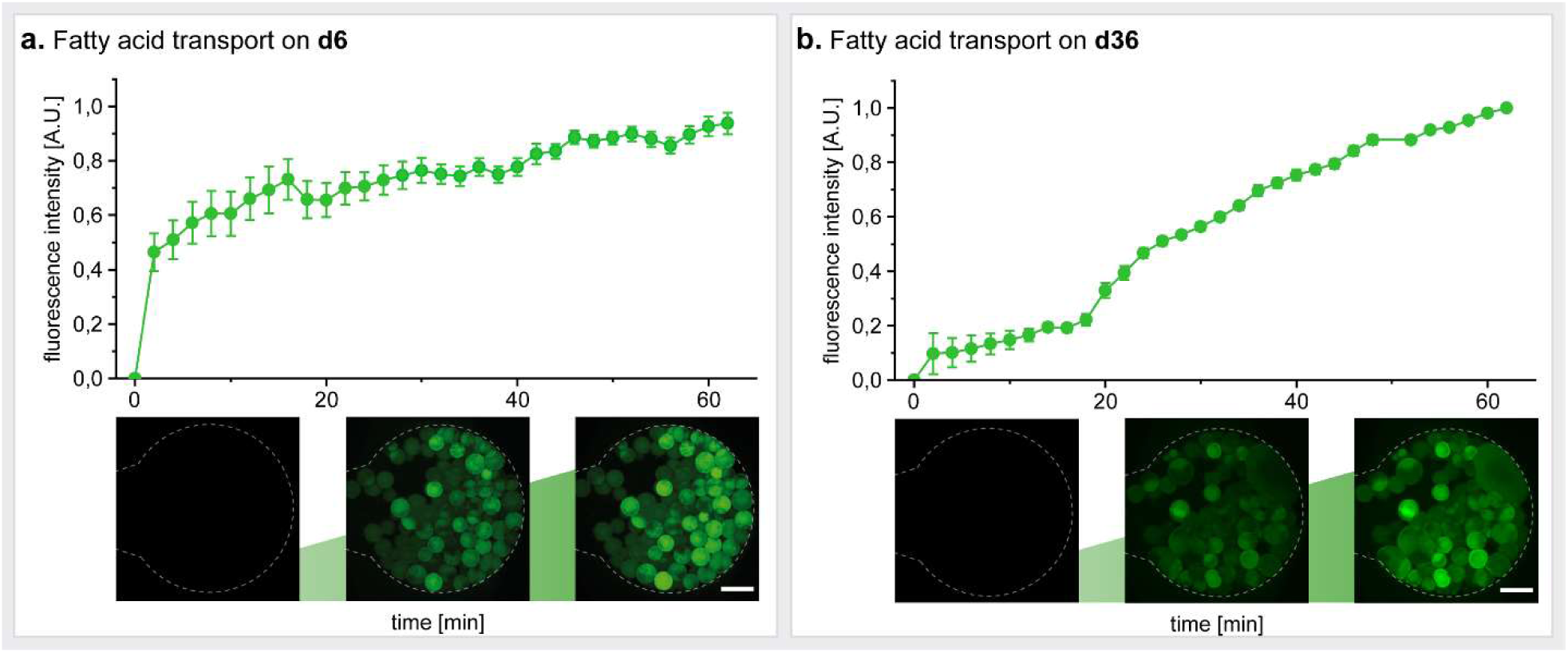
Long-term functionality of the human WAT-on-a-chip. Dynamic measurement of the average fluorescence intensity in tissue chambers perfused with medium containing a fluorescent fatty acid analog (4 µM BODIPY™ 500/510 C1, C12) revealed functional fatty acid uptake and accumulation in WAT after (**a.**) 6 as well as (**b.**) 36 days of on-chip culture (n = 6 individual tissue chambers). Scale bars equal 150 µm.

### Applicability of WAT-on-a-chip for drug testing

Since an important area of application for the organ-on-a-chip technology is drug development, we conducted a proof of concept compound test to assess the applicability of our WAT-on-a-chip for pharmaceutical research. After 6 days of on-chip culture, we exposed the WAT to the β-adrenergic agonist, isoproterenol, which is known to induce lipolysis. By additionally supplementing the medium with the fluorescently tagged fatty acid analog (BODIPY™ 500/510 C1, C12) for 60 minutes, we were able to monitor isoproterenol-related effects on fatty acid uptake and release by WAT, using standard fluorescence microscopy (**figure 6**). While media with fluorescently tagged fatty acids was perfused, the normalized fluorescent intensity in the isoproterenol-treated systems increased significantly more slowly than in non-treated control systems. After switching to media without the fluorescently tagged fatty acid analog, the fluorescent intensity decreased much quicker in the isoproterenol-treated systems. Taken together, this means that the isoproterenol exposure induced a reduction of the net uptake rate of fatty acids and an increase of their release rate. Both findings are in line with the expected lipolysis-inducing effect of isoproterenol.

**Fig. 6.**
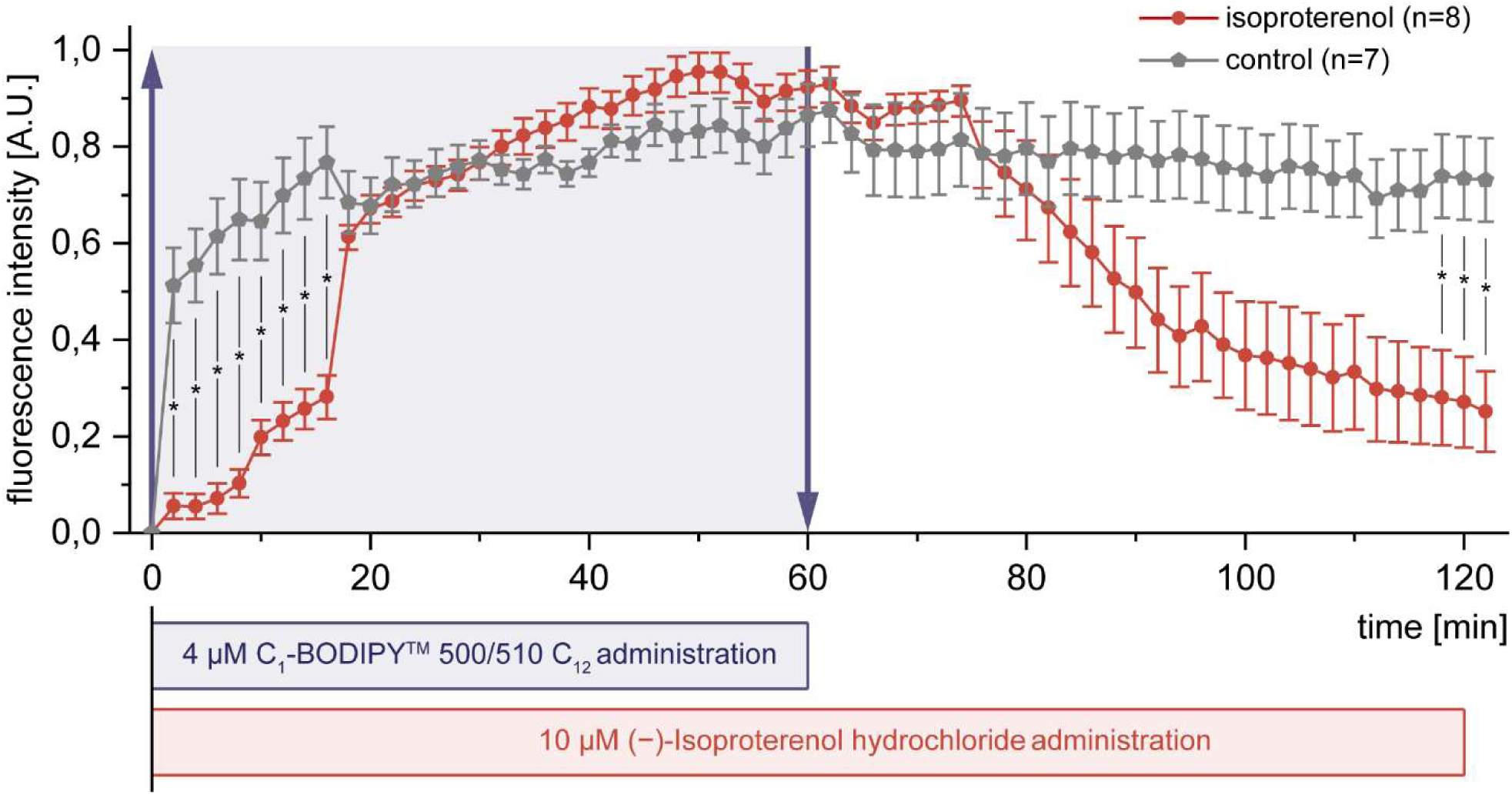
Applicability for drug testing illustrated by the effect of isoproterenol on fatty acid uptake and release. Normalized mean fluorescence intensity obtained by time-lapse fluorescence microscopy of WAT-on-a-chip systems perfused with medium supplemented with the fluorescently tagged fatty acid analog (4 µM BODIPY™ 500/510 C1, C12) during the first 60 minutes. Kinetics in systems exposed to 10 µM Isoproterenol (red circles) show significant differences to non-treated controls (grey polygons) (n = 7 and n = 8 individual tissue chambers; mean fluorescence intensities ± SEM are presented).

Evidently, the used approach based on standard microscopy provides facile and non-invasive read-outs for drug screening, with a very high time-resolution, which is amenable for massive parallelization and automation. One noteworthy limitation is the requirement for normalization of readings from individual tissue chambers. Absolute quantitative values would provide even more information and potential for cross-correlation; it is, therefore, planned to address this in future generations of chips and readout-infrastructure. Overall, the successful proof of concept highlights the applicability of the WAT-on-a-chip system for screening purposes and the potential the system can have for drug development purposes.

## Conclusion

Due to the lack of suitable human model systems, research in adipose tissue biology and obesity has so far relied mostly on animal models, non-physiological *in vitro* systems, GWAS studies and (clinical) epidemiology. The first two approaches have major limitations in terms of their translatability to humans; the latter two require complex statistical analyses of large data sets and do not permit strong conclusions on pathophysiological mechanisms or efficacy of therapeutic interventions in individuals. The human WAT-on-a-chip platform presented here provides a novel tool that enables the maintenance, monitoring and manipulation of human adipose tissue in a durably stable, microphysiological environment. The vasculature-like perfusion provides opportunities for biomarker evaluation and “liquid biopsies” that can be cross-correlated with clinical endpoints, as well as a potential connection link for integration with other organ-on-a-chip platforms.(32) Multi-organ-chips integrating WAT are likely to be of major interest e.g. for ADMET, diabetes or non-alcoholic steatohepatitis (NASH) studies.

Overall, the WAT-on-a-chip system presented here, offers new perspectives and opportunities in mechanistic studies, pharmaceutical development and testing, as well as in personalized medicine.

## Materials and Methods

### Fabrication and characterization of microfluidic platforms

#### Chip fabrication by soft lithography and replica molding

The microfluidic platform is a custom-designed three-layered hybrid device featuring two micro-patterned polydimethylsiloxane (PDMS; Sylgard 184, Dow Corning, USA) layers, which are separated by an isoporous semipermeable membrane. The media channel and tissue chamber microstructures in the PDMS slabs were generated using two differently patterned master wafers that served as positive molding templates. The intricately structured masters were fabricated by commonly used photolithographic processes described previously.(34) The chips’ PDMS structures were created using two different molding techniques: standard molding to obtain thicker slabs with closed structures, and exclusion molding to obtain thin layers with open structures (cf. “Replica molding of PDMS parts” in the supplements section). To prepare the semipermeable membranes, commercially available polyethylene terephthalate (PET) membranes (r_P_ =3 µm; ρ_P_ = 8 × 10^5^ pores per cm^2^; TRAKETCH® PET 3.0 p S210×300, SABEU GmbH & Co. KG, Northeim, Germany) were functionalized by a plasma-enhanced, chemical vapor deposition (PECVD) process (cf. “Membrane functionalization” in the supplements section). In a final step, chips were assembled in three subsequent O_2_-plasma activation (15 s, 50 W; Diener Zepto, Diener electronic GmbH + Co. KG, Ebhausen, Germany) and bonding steps, followed by an overnight exposure to 60 °C for bonding enhancement (cf. “Chip assembly” in the supplements section).

#### Chip preparation for experiments

On the day before cell injection, chips were O_2_-plasma sterilized (60 s, 50 W) and subsequently filled with Dulbecco’s phosphate-buffered saline without MgCl_2_ and CaCl_2_ (PBS^-^; Sigma-Aldrich Chemie GmbH, Steinheim, Germany), under sterile conditions. Then, the PBS^-^ filled chips were kept overnight at 4 °C in PBS^-^ to allow evacuation of residual air from the channel systems.

#### Numerical modeling

To model fluid flow and transport of a diluted species, COMSOL Multiphysics (COMSOL, Stockholm, Sweden) software was used. The process was based on a numerical model which was previously published for our previous WAT-on-a-chip(34). Briefly, the incompressible stationary free fluid flow was modeled by the Navier-Stokes equation with the properties of water (dynamic viscosity μ = 1 x 10^-3^ m^2^/s, density ρ = 1000 kg/m^3^) at a flow of 20 µl/h. Fluid flow from the media channel through the isoporous membrane into the tissue channel was modeled using Darcy’s law (porosity = 0.056, hydraulic permeability κ = 1.45 x 10-14 m^2^). The transport of diluted species was described by the time-dependent convection-diffusion with a diffusion coefficient 1 x 10^-9^ m^2^/s and an initial concentration of 1 mol/m^3^.

#### Diffusive transport

To visualize diffusion of compounds from the media microchannels over the isoporous membrane into the tissue chamber inside the microfluidic platform, we monitored the perfusion of a 0.5 mg/ml fluorescein isothiocyanate–dextran (FITC-dextran; 150 kDa, 46946, Sigma-Aldrich Chemie GmbH) solution in PBS^-^ in the system. Prior to the diffusion experiment, the tissue chambers were filled with the collagen hydrogel matrix commonly used to encapsulate the adipocytes (cf. “Injection of human mature adipocytes into the microfluidic platform”). The flow rate was set to 40 µl/h and the fluorescence intensity was measured every 6.2 s for three positions in the chip – the media channel, the top of the underlying tissue chamber, as well as the bottom of the same tissue chamber.

### Isolation and culture of primary human adipocytes

#### Human tissue samples

Human adipose-tissue biopsies were obtained from plastic surgeries performed by Dr. Ulrich E. Ziegler (Klinik Charlottenhaus, Stuttgart, Germany). All procedures were carried out in accordance with the rules for medical research of human subjects, as defined in the Declaration of Helsinki. Patients signed a written consent form according to the Landesärztekammer Baden-Württemberg (IRB# F-2012-078).

All primary mature adipocytes were isolated from biopsies that were taken from female, pre-obese donors (BMI 25.0 - 29.9, as per the WHO classification), aged 45 to 55.

#### Isolation of primary human adipocytes

Primary mature adipocytes were isolated from subcutaneous adipose tissue samples. The isolation protocol was performed as previously described(44,45), with slight modifications. In brief, the subcutaneous adipose tissue was rinsed twice with Dulbecco’s phosphate buffered saline with MgCl_2_ and CaCl_2_ (PBS^+^; Sigma-Aldrich Chemie GmbH), and visible blood vessels, as well as connective-tissue structures, were thoroughly removed. The remaining adipose tissue was cut into fine pieces of approximately 1 cm^3^, and digested with a collagenase solution [(0.13 U/ml collagenase type NB4 (Serva Electrophoresis GmbH, Heidelberg, Germany) in Dulbecco’s Modified Eagle Medium (DMEM/Ham’s-F12; Thermo Fisher Scientific, Waltham, USA), with 1% bovine serum albumin (BSA; Sigma-Aldrich Chemie GmbH)] for 60 min at 37 °C on a rocking shaker (50 cycles/min; Polymax 1040, Heidolph Instruments GmbH & CO. KG, Schwabach, Germany). Next, the digested tissue was passed through cell strainers (mesh size: 500 µm), and subsequently washed three times with DMEM/Ham’s-F12. For each washing step, adipocytes and medium were mixed, and left to rest for 10 min to allow separation of the buoyant adipocytes and the medium; this was followed by aspiration of the liquid media from underneath the packed layer of adipocytes.

#### Injection of human mature adipocytes into the microfluidic platform

Immediately after the isolation of white adipocytes from tissue samples, the adipocytes were prepared for injection into the microfluidic platforms. Sixty µl of densely packed adipocytes were mixed with 24 µl of a dispersion of 10 mg/ml collagen type 1 (from rat tail, provided by Fraunhofer IGB), 6 µl neutralization buffer [DMEM/Ham’s F-12 (10X); Biochrom GmbH, Berlin, Germany), and 50 mM NaOH in Aqua dest (1:1) with 0.2 M NaHCO_3_ and 0.225 M HEPES (Carl Roth GmbH + Co. KG, Karlsruhe, Germany)] and immediately injected into the chip’s tissue-chamber system by manual pressure. Each system of connected tissue chambers was loaded individually at a steady pace, to ensure that the collagen hydrogel reached the tissue chambers before the onset of gelation.

#### On-chip culture of adipose tissue

During on-chip culturing, the WAT-chips were maintained in a humidified incubator at 37°C and a 5% CO_2_ atmosphere. The adipocytes were supplied with a 20-40 µl/h flow of Subcutaneous Adipocyte Maintenance Medium (AM-1; BioCat GmbH, Heidelberg, Germany) maintained by positive pressure from an automated syringe pump (LA-190, Landgraf Laborsysteme HLL GmbH, Langenhagen, Germany). Under sterile conditions, media reservoirs were re-filled with fresh media every other day, and the media effluents were collected from the media-systems’ outlets once every 1-2 days (depending on the experiment), and stored at −80°C for subsequent analysis of metabolites.

### Structural and functional characterization of on-chip adipose tissues

#### Fluorescent double staining of intracellular lipid vacuoles and nuclei

To visualize the structure of on-chip adipose tissues, intracellular lipid vacuoles and nuclei were stained using the neutral lipid stain BODIPY™ 493/503 (4,4-Difluoro-1,3,5,7,8-Pentamethyl-4-Bora-3a,4a-Diaza-s-Indacene; D3922, Thermo Fisher Scientific) and 4′,6-Diamidin-2-phenylindol (DAPI;D8417, Sigma-Aldrich Chemie GmbH). All fixation-and staining solutions were flushed through the chip at a rate of 80 µl/h by a syringe pump. The in-chip adipocytes were fixed overnight with 4% phosphate-buffered formaldehyde solution (Roti®-Histofix 4 %, P087, Carl Roth GmbH + Co. KG). Next, permeabilization was achieved by flushing the chip for 3 h with PBST [PBS^+^ with 0.1% Tween-20 (P7949, Sigma-Aldrich Chemie GmbH)]. Then, the fluorescence staining solution – PBST with 1 µg/ml BODIPY™ 493/503 and 1 µg/ml DAPI – was pumped through the system for 2 h, and finally washed out with PBS^+^ for at least 30 min to remove residual staining solution. Imaging of the stained adipocytes was performed and processed by a laser scanning microscope (Zeiss LSM 710, Carl Zeiss, Oberkochen, Germany) with specialized software (ZEN 2012 SP1 (black edition), Release Version 8.1)).

#### Live/dead staining of integrated white adipose tissue

To assess the viability of the on-chip WAT, a live/dead-assay based on fluorescein diacetate (FDA; F1303, Thermo Fisher Scientific) and propidium iodide (PI; P3566, Thermo Fisher Scientific) was performed. Prior to staining, the culture medium was removed from the chip by flushing the media channels with PBS, using gravitational flow. The staining solution containing FDA (final concentration 27 µg/ml) and PI (final concentration 135 µg/ml) diluted in PBS, was flushed into the media channel by gravitational flow, and incubated for 25 min in a humidified incubator at 37°C and 5% CO_2_. The staining solution was then removed by flushing PBS through the media channels using gravitational flow. Right after that, imaging was performed by an inverted fluorescence microscope (Leica DMi8, LEICA Microsystems GmbH, Wetzlar, Germany).

#### Monitoring and analysis of fatty acid uptake and release

For online monitoring of fatty acid uptake by the WAT, 4 µM of the fluorescently-labeled fatty acid analog, BODIPY™ 500/510 C1, C12 (4,4-Difluoro-5-Methyl-4-Bora-3a,4a-Diaza-s-Indacene-3-Dodecanoic Acid;D3823, Thermo Fisher Scientific), was added to the culture medium for a duration of 60 min and pumped through the media systems at 80 µl/h via positive pressure provided by a syringe pump. Next, the culture medium was switched to AM-1 only, to visualize the release of the fluorescent fatty acid analog. During the experiment, the chips were placed under complete darkness in an incubator chamber that was fitted to the microscope stage, and set to 37 °C. Imaging was performed using the inverted Leica DMi8 fluorescence microscope. During the 120 min running time of the experiment, images of both bright-field (BF) and FITC-channels were captured every two minutes from each tissue chamber on the chip. As a reference, each position was imaged at time point 0 min (t0).

To quantify the patterns of fluorescence intensity during the uptake and release of the fluorescent fatty acid analog, the mean gray value of the fluorescence of the individual tissue chambers and the fluorescence of the background were measured for each time point using ImageJ 1.50i software (National Institute of Health, Bethesda, USA). After subtracting the background levels from the fluorescence intensities measured in the chambers, the offset was calculated by setting the intensity measured at t0 to a fluorescence intensity of 0 A.U. Then, the intensities for each chamber were expressed as a percentage of the highest recorded intensity from that chamber during the experiment. Note: Normalization on the chamber-level was considered to be necessary, because the amount of injected adipocytes varied between the chambers with the used WAT-chip design.

#### Responsiveness to β-adrenoreceptor agonistic drugs

To assess the effect of β-adrenergic agonist drugs on fatty acid metabolism, AM-1 medium was supplemented with 10 µM isoproterenol hydrochloride (I6504, Sigma-Aldrich Chemie GmbH). Fluorescence intensity was measured for 60 min with, and then for 60 min without addition of 4 µM BODIPY™ 500/510 C1, C12 to the conditioned medium at 37°C with an inverted fluorescence microscope (LEICA DMi8, LEICA Microsystems GmbH) to assess the dynamics of fatty acid uptake and release. Measurement and analysis of fluorescence was performed as described above.

#### Analysis of adipose-associated metabolites

The non-esterified fatty acid (NEFA) contents of medium effluents were determined by enzymatic analysis, using the ACS-ACOD-MEHA method (NEFA-HR(2) Assay, FUJIFILM Wako Chemicals Europe GmbH, Neuss, Germany). Effluents were thawed and centrifuged at 10,000 x g for 10 min. Then, 25 µl of sample [i.e. the effluents’ supernatant, or AM-1 as blank, or oleic acid for a standard curve (270-77000, FUJIFILM Wako Chemicals Europe GmbH)] were supplemented with 100 µl of the R1 solution (434-91795, FUJIFILM Wako Chemicals Europe GmbH) and 50 µl of the R2 solution (436-91995, Wako Chemicals GmbH), incubating 10 min at 37°C after each addition. NEFA concentrations were determined by measuring absorbance at 550 nm (Infinite® 200 PRO, Tecan Trading AG, Switzerland). To assess the influence of albumin on NEFA release, 0.2% human serum albumin (HSA; A1653, Sigma-Aldrich Chemie GmbH) was added to the culture medium for 24 h. To ensure that increases in the measured NEFA concentrations could not be attributed to the presence of HSA during assay performance, we performed NEFA-measurements on the oleic acid standard with and without HSA. No differences were observed in the detected NEFA-concentration between the two conditions (**figure S4**).

As another readout for lipolysis, glycerol concentrations were determined using a colorimetric assay. Media effluents were thawed and centrifuged at 10,000 x g for 10 min. Then, 40 µl of sample [i.e. the effluents’ supernatants, or AM-1 as blank, or Glycerol Standard Solution for a standard curve (G7793, Sigma-Aldrich Chemie GmbH)] were supplemented with 60 µl of the Free Glycerol Reagent (F6428, Sigma-Aldrich Chemie GmbH). After a 5-min incubation at 37°C, absorbance at 540 nm was recorded.

#### Statistical analysis

All graphs show raw data means ± standard deviation (unless otherwise indicated). We annotated or sample size n as follows:

- For analyses pertaining to data at tissue chamber level (e.g. from fatty acid transport assays), n denotes the number of tested tissue chambers.
- For analyses pertaining on tissue system level only, i.e., the collectivity of eight chambers connected via one media channel, (e.g. metabolite measurements), n denotes the number of tested tissue systems.

For statistical analysis of differences, we performed independent two-sample **t**-tests using OriginPro 2018 software (OriginLab, Northhampton, MA, USA). Unless otherwise indicated, a **p**-value threshold for significance of 0.05 was used.

## Acknowledgements

The authors thank Dr. Ziegler (Klinik Charlottenhaus, Stuttgart) for the kind provision of human adipose tissue from plastic surgery, Dr. Jakob Barz for help with the PECVD coating process, and Silvia Kolbus-Hernandez, Elena Rubiu as well as Luisa Merz for their assistance with cell culture and chip fabrication. We further thank Prof. Andreas Stahl and Pete Zushin for helpful discussions and Joost Overduin for language-editing. The research was supported in part by the Fraunhofer-Gesellschaft internal programs Talenta Start (to J.R.) and Attract (601543), the DAAD funded by the Bundesministeriums für Bildung und Forschung (BMBF) (PPP USA 2018, 57387214) (all to P.L.), as well as the Ministry of Science, Research and the Arts of Baden-Württemberg (Az: 7542.2-501-1/13/6 to P.L. and Az: 33-729.55-3/214-8 to K.S.-L.).

## Author Contributions

J.R. and P.L. designed the device. J.R. and C.B. fabricated the device. J.R., C.B. and E.K. performed and analyzed the experiments. S.S. performed membrane characterization. C.P. performed simulations. K.S.-L. gave biological advice. J.R., K.S.-L. and P.L. wrote the manuscript. J.R. and P.L. designed the study.

## Competing interests

None

## Supplements

### Microfabrication of the microfluidic platform

#### Replica molding of PDMS parts

Polydimethylsiloxane (PDMS; Sylgard 184, Dow Corning, USA) layers were created *via* replica molding (cf. **figure S1**) of a 10:1 (w/w) mixture of PDMS pre-polymer and curing agent. To create the PDMS layer containing the media channel structures by standard molding, 25 g of the uncured PDMS mix was poured on top of the wafer and stored overnight at 60 °C for curing of the PDMS. On the next day, after peeling off the standard-molded slab of PDMS from the media wafer, inlets and outlets were created using a biopsy punch (diameter: 0.75 mm; World Precision Instruments). In contrast, an exclusion molding technique was used to create the PDMS layer incorporating the open tissue-chamber structures: ∼ 1 g of the uncured PDMS/curing agent mixture was poured on top of the micropatterned wafer and spread by gently tilting and rotating the wafer. A pre-cut, rounded piece of 3M Scotchpak™ 1022 Release Liner Fluoropolymer Coated Polyester Film (3M ID 70000200280, St. Paul, MN, USA) was lowered onto the PDMS mixture, with its treated surface face-down. The ensemble was clamped between to smooth surfaces (e.g. unpatterned wafers), and stored overnight at 60 °C for curing of the PDMS. The exclusion-molded PDMS structures were carefully examined, and voided of residues of PDMS adhering to the foil, that might close off the open structures.

**Fig S1:**
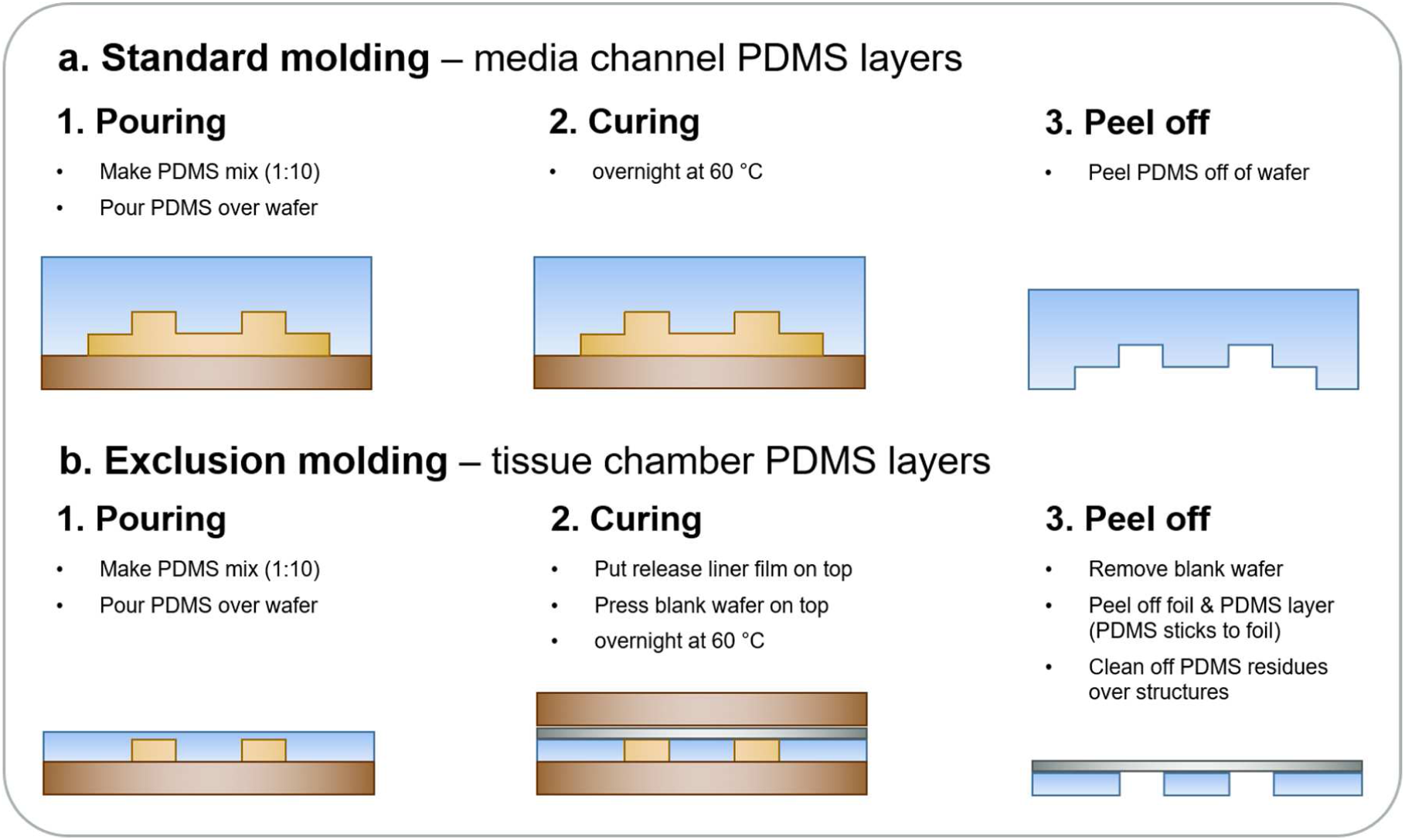
Molding techniques for fabricating the WAT-chip. **a.** PDMS slabs featuring media channel microstructures were produced by a standard molding process. First, 25 g of a PDMS mixture (containing a 10:1 w/w ratio of PDMS pre-polymer and curing agent) was poured over the media wafer. Then, after curing overnight at 60 °C, the PDMS mold was peeled off the wafer. **b.** PDMS layers featuring the tissue chamber systems were fabricated by exclusion molding. For this procedure, only ∼ 1 g of the PDMS mixture was poured over the microstructured wafer. Then, a release liner, consisting of a polyester film, coated with fluoropolymer, was placed on top of the PDMS mixture, with the coated side facing the PDMS. A blank wafer was pressed against the other side of the foil to create open PDMS structures. After peel-off, PDMS stuck to the foil and PDMS residues were gently removed to uncover the intended open structures.

#### Membrane functionalization

Commercially available PET-membranes (TRAKETCH® PET 3.0 p S210×300, SABEU GmbH & Co. KG, Northeim, Germany) were modified using a plasma-enhanced chemical vapor deposition (PECVD) process. The PECVD, carried out by a low-pressure plasma, was performed in a custom-built plate reactor. The plasma was generated using a high-frequency (HF) source at 13,56 MHz (CESAR™ Generator Model 136, Dressler Hochfrequenztechnik GmbH, Stolberg, Germany) and a matching box (AE VarioMatch™ Match Network, VM 1500 Platform, Dressler Hochfrequenztechnik GmbH, Stolberg, Germany). The PECVD reactor was evacuated by a rotary vane vacuum pump (TRIVAC D 40B, Leybold GmbH, Köln, Germany) and a roots vacuum pump (RUVAC WS 251, Leybold GmbH, Köln, Germany). Pressure measurements were carried out using a vacuum gauge controller (Granville-Phillips® Series 375 Convectron®, MKS Instruments, Andover, MA, United States). All gas flows into the reactor were controlled by mass-flow controllers (Type 1259, MKS Instruments, Andover, MA, United States) and a multi-gas controller (Type 647B, MKS Instruments, Andover, MA, United States). The precursor Hexamethyldisiloxane (HMDSO) was evaporated at 37 °C and led into the reactor controlled by a mass-flow controller (UFC-7300, UNIT Instruments Inc., Austin, TX, United States). Before initiation of the coating process, the reactor chamber was cleaned using isopropanol, and all gas lines were evacuated to remove any residual gas. The PET-membranes, sized 210 × 300 mm^2^, were placed inside the reactor chamber on top of a piece of protection paper of the same size, provided by the supplier. The plasma reactor was closed and evacuated until a pressure of approximately 1 - 2 µbar was reached. Before starting the PECVD, the membranes were pre-treated by a 30-s exposure to an oxygen plasma to clean and activate the substrate. The process parameters are listed in table S1. Immediately after the pre-treatment, the membranes were coated by PECVD for 40 sec, using the parameters in table S1. The pre-treatment and coating process were applied on both sides of the membranes. After the membrane functionalization, the membranes were cut to the desired size, using a CO_2_-laser-cutter (VLS2.30, Universal Laser Systems).

**Table S1:**
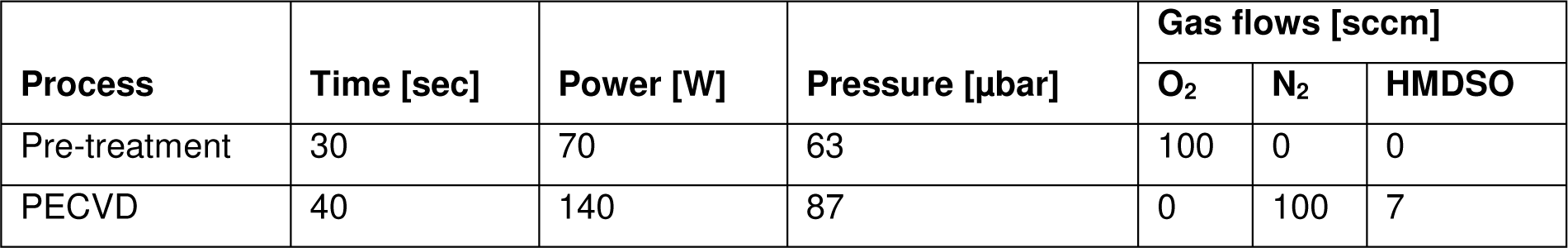
PECVD process parameters

| Process | Time [sec] | Power [W] | Pressure [µbar] | Gas flows [sccm] |  |  |
| --- | --- | --- | --- | --- | --- | --- |
|  |  |  |  | O<sub>2</sub> | N<sub>2</sub> | HMDSO |
| Pre-treatment | 30 | 70 | 63 | 100 | 0 | 0 |
| PECVD | 40 | 140 | 87 | 0 | 100 | 7 |

### Adhesion test of PET to PDMS bonding

Peel tests were carried out to assess the bonding strength between the functionalized membranes and the PDMS. Membranes were lasercut in pieces sized 115 × 32 mm^2^. After washing with 70 % ethanol, drying and fixing to a 175 µm thick PET foil, membranes and PDMS were plasma activated and bonded, as described above. After the completion of bonding, 180° peel tests (width B = 32 mm) were conducted using a tensile testing machine (Z005; Zwick Roell, Ulm, Germany) and a 2,5 kN nominal force load cell (XForce HP, Zwick Roell, Ulm, Germany). To test the longtime stability of bonding, membranes bonded to PDMS were kept in PBS-for 7 days at room temperature.

The membranes stored for 7 days in PBS^-^ and membranes tested straight after bonding were compared (figure S2) and no weakening of the bonding was observed. In contrast to conventional membrane bonding approaches, this PECVD-based bonding technique allows for a facile generation of PDMS-PET hybrids that is easily scalable.

**Fig S2:**
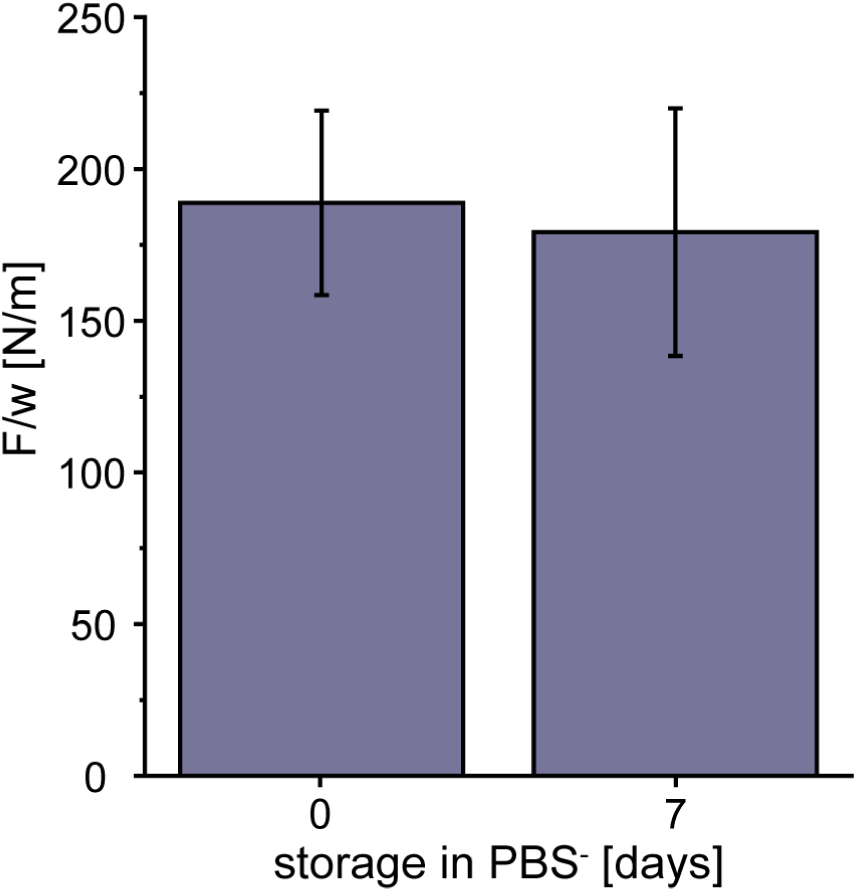
Adhesion test of PET to PDMS bonding. As confirmed by 180° peel tests, the bonding strength between PECVD-funtionalized PET membranes and PDMS was not compromised by long-term storage in PBS^-^.

#### Chip assembly

Prior to chip assembly, all PDMS parts were thoroughly rinsed with isopropanol, blow-dried with N_2_, and further cleaned by repeated pressing and peeling with strips of household adhesive tape. In a three-step process (cf. figure S3) of O_2_-plasma activation (15 s, 50 W; Diener Zepto, Diener electronic GmbH + Co. KG, Ebhausen, Germany) and bonding, the exclusion-molded layer was bonded to a microscope slide (bonding enhancement for 30 min at 60°C) (1), the functionalized PET-membrane was bonded into the insert within the lop layer (2), and finally, the top layer and membrane were bonded to the bottom layer on the glass slide (3). Alignment of structures in top and bottom layer was conducted under visual guidance of a stereomicroscope. To enhance bonding, the chip was firmly pressed together between two microscope slides using foldback clips and kept overnight in an oven at 60°C. To verify the sealing tightness and fluidic connectivity of the system, tissue chambers and media channels were perfused with 70% ethanol.

**Fig S3:**
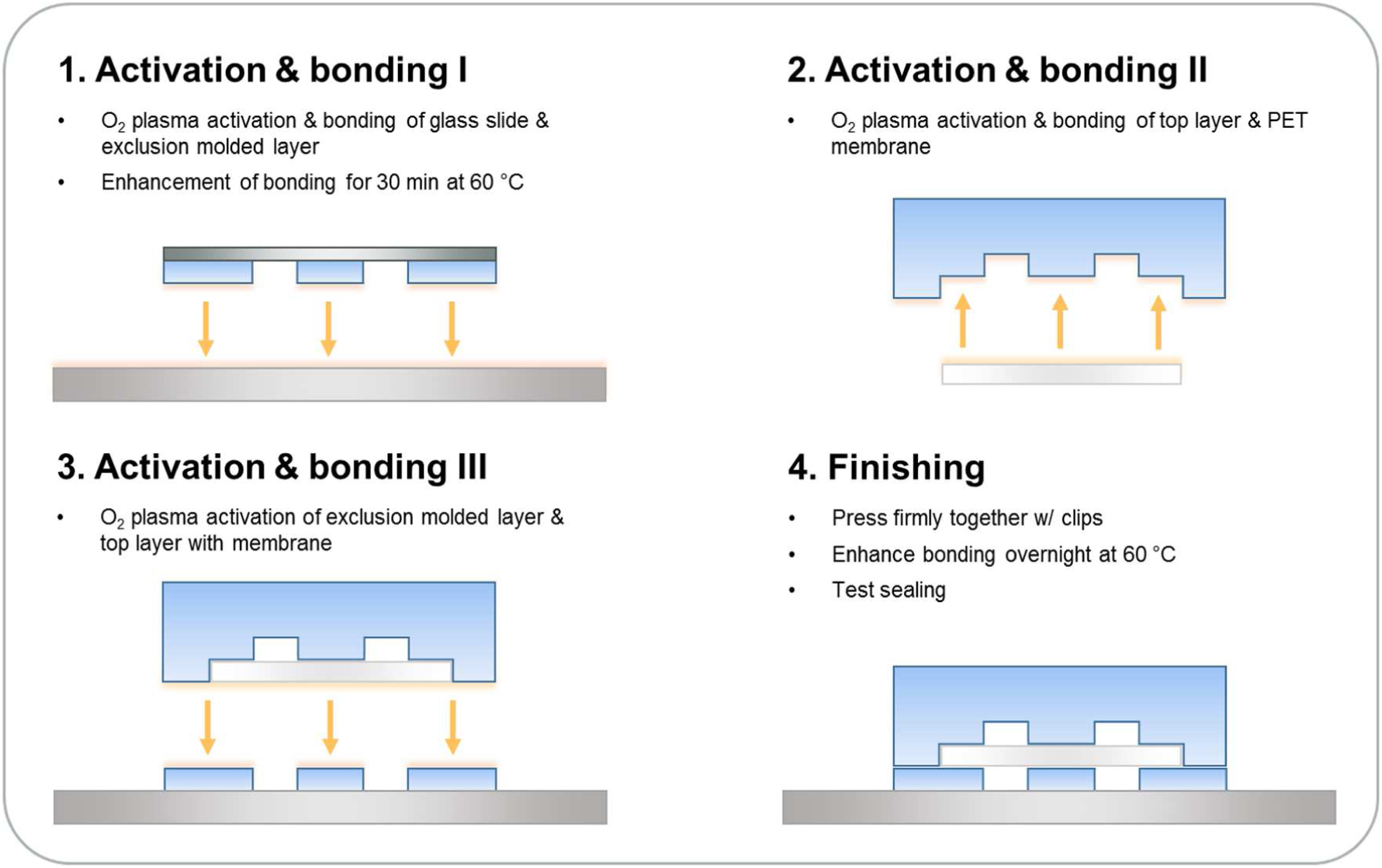
Chip assembly supported by plasma activation. The WAT-chip was assembled in three subsequent O_2_-plasma activation (15 s, 50 W) and bonding steps: 1. Bonding of the exclusion-molded layer to a microscope slide and enhancement of bonding for 30 min at 60 °C. 2. Bonding of the PET membrane to the standard-molded PDMS layer. 3. Bonding of the top layer to the exclusion-molded layer. 4. Overnight enhancement of bonding at 60 °C. To test sealing strength, and fluidic continuity, 70% ethanol was flushed through the systems.

#### Influence of human serum albumin presence on non-esterified fatty acid analysis

To ensure that increases in the measured NEFA concentrations could not be attributed to the presence of HSA during assay performance, we performed NEFA-measurements of the oleic acid standard with and without HSA supplemented medium. Oleic acid standard (100 µM) was added to AM-1 only or to AM-1 supplemented with 0.2% (w/v) HSA, and the NEFA assay was performed as described. No differences were observed in the measured absorbances between the two conditions (**figure S4**).

**Fig S4:**
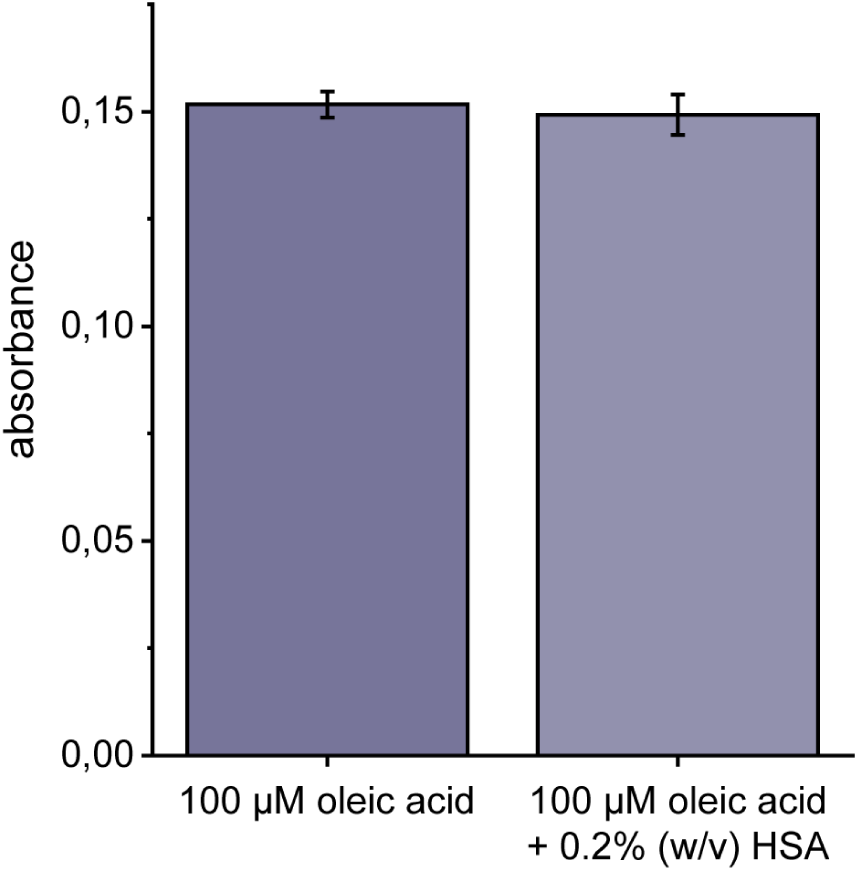
Influence of HSA presence on NEFA measurement. The presence of 0.2% (w/v) HSA in the culture medium did not influence the enzymatic analysis of the NEFAs.

